# Dengue fever and *Aedes aegypti* risk in the Galápagos Islands, Ecuador

**DOI:** 10.1101/113829

**Authors:** Ryan Nightingale, Catherine Lippi, Sadie J. Ryan, Mercy J. Borbor-Cordova, Marilyn Cruz B, Fernando Ortega, Renato León, Egan Waggoner, Anna M. Stewart Ibarra

**Affiliations:** Department of Medicine, SUNY Upstate Medical University, Syracuse, NY, USA; Center for Global Health and Translational Science, SUNY Upstate Medical University, Syracuse, NY, USA; Department of Geography, University of Florida, Gainesville, FL, USA; Emerging Pathogens Institute, University of Florida, Gainesville, FL, USA; College of Life Sciences, University of KwaZulu Natal, Durban, South Africa; Escuela Superior Politécnica del Litoral, Guayaquil, Guayas, Ecuador; Agencia de Regulación y Control de la Bioseguridad y Cuarentena para Galápagos (ABG), Puerto Ayora, Galápagos, Ecuador; School of Public Health, Universidad San Francisco de Quito, Quito, Pichincha, Ecuador; Laboratorio de Entomología Médica & Medicina Tropical, LEMMT, Universidad San Francisco de Quito, Quito, Pichincha, Ecuador

**Keywords:** dengue fever, Aedes aegypti, social-ecological risk, islands, Galápagos, Ecuador

## Abstract

**Introduction:** Dengue fever is an emerging infectious disease in the Galápagos Islands of Ecuador, with the first cases reported in 2002 and periodic outbreaks since then. Here we report the results of a pilot study conducted in two cities in 2014: Puerto Ayora (PA) on Santa Cruz Island, and Puerto Baquerizo Moreno (PB) on Santa Cristobal Island. The aims of this study were to assess the social-ecological risk factors associated with dengue and mosquito presence at the household-level.

**Methods:** In 2014 we conducted 100 household surveys (50 on each island) in neighborhoods with prior reported dengue. Adult mosquitoes were collected inside and outside the home, larval indices were determined through container surveys, and heads of households were interviewed to determine demographics, prior dengue infections, housing conditions, and knowledge, attitudes and practices regarding dengue. Multimodel selection methods were used to derive best-fit generalized linear regression (GLM) models of prior dengue infection, and the presence of *Ae. aegypti* in the home.

**Results:** We found that 24% of PB and 14% of PA respondents self-reported a prior dengue infection, and more PB homes than PA homes had *Ae. aegypti.* The top-ranked model for prior dengue infection included human movement – travel between neighborhoods, between islands, and to the mainland; demographics including salary level and education of the head of household, and increase with more people per room in a house, house condition, access to water quality issues, and dengue awareness. The top-ranked model for the presence of *Ae. aegypti* included housing conditions, including the presence of window screens and air conditioners, mosquito control actions, and dengue risk perception.

**Discussion/conclusion:** To our knowledge, this is the first study of dengue risk and *Aedes aegypti* in the Galápagos Islands. The findings that human movement within and between islands, and to and from the mainland, were important to reported dengue cases confirms concerns of this route of introduction and repeated transmission.

## Background

Dengue fever is a mosquito-borne viral illness that has rapidly increased in geographic distribution and incidence in recent decades [1]. In the early 2000s, dengue began to emerge in relatively isolated islands, including the Galápagos Islands of Ecuador [2], the Hawaiian Island of the United States [3], and Easter Islands of Chile [4-6]. Islands can be viewed as valuable sources of epidemiological research due to their relative isolation, which makes them ideal for measuring the response of dengue transmission to local factors without regional influences [7].

The dengue virus (DENV, family *Flaviviridae,* genus *Flavivirus*) causes an estimated 96 million new apparent (symptomatic) infections per year worldwide, with 16 million infections in the Americas [8]. In recent years, new arboviruses transmitted by the same mosquito vectors (*Aedes aegypti* and *Aedes albopictus*) have emerged the Americas, including chikungunya virus and zika virus. Mosquito control by the public health sector is the primary means of disease control intervention available; however, existing vector control efforts have been largely unsuccessful at preventing epidemics. New strategies that consider the nuances of local social-ecological conditions and risk factors are urgently needed to inform local targeted control campaigns, and to understand effective household-level strategies and behaviors.

The Galápagos Islands of Ecuador are a World Heritage Site, located 1000 kilometers from the mainland (Figure 1). The islands are renowned for their biodiversity due to their relative isolation and minimal impact by people, who first settled on the islands in the mid 1800s [9]. In recent decades, the population has increased rapidly, from 1,346 people in 1950 [9] to 30,890 people residing on four islands in 2017 [10]. Tourism is the most important economic activity [9], with 215,691 tourists visiting the islands in 2014 [11]. Invasive species introduced due to human activity, including pathogens, have been recognized as a serious threat to endemic species on the islands [12]; however, less attention has been paid to emerging pathogens in humans. Prior studies of West Nile Virus risk on the Galápagos focused on the impacts on endemic bird species [13–15]. However, emerging infections in humans are a growing concern due to the increasingly urban resident population and the large number of international tourists.

**Figure 1.**
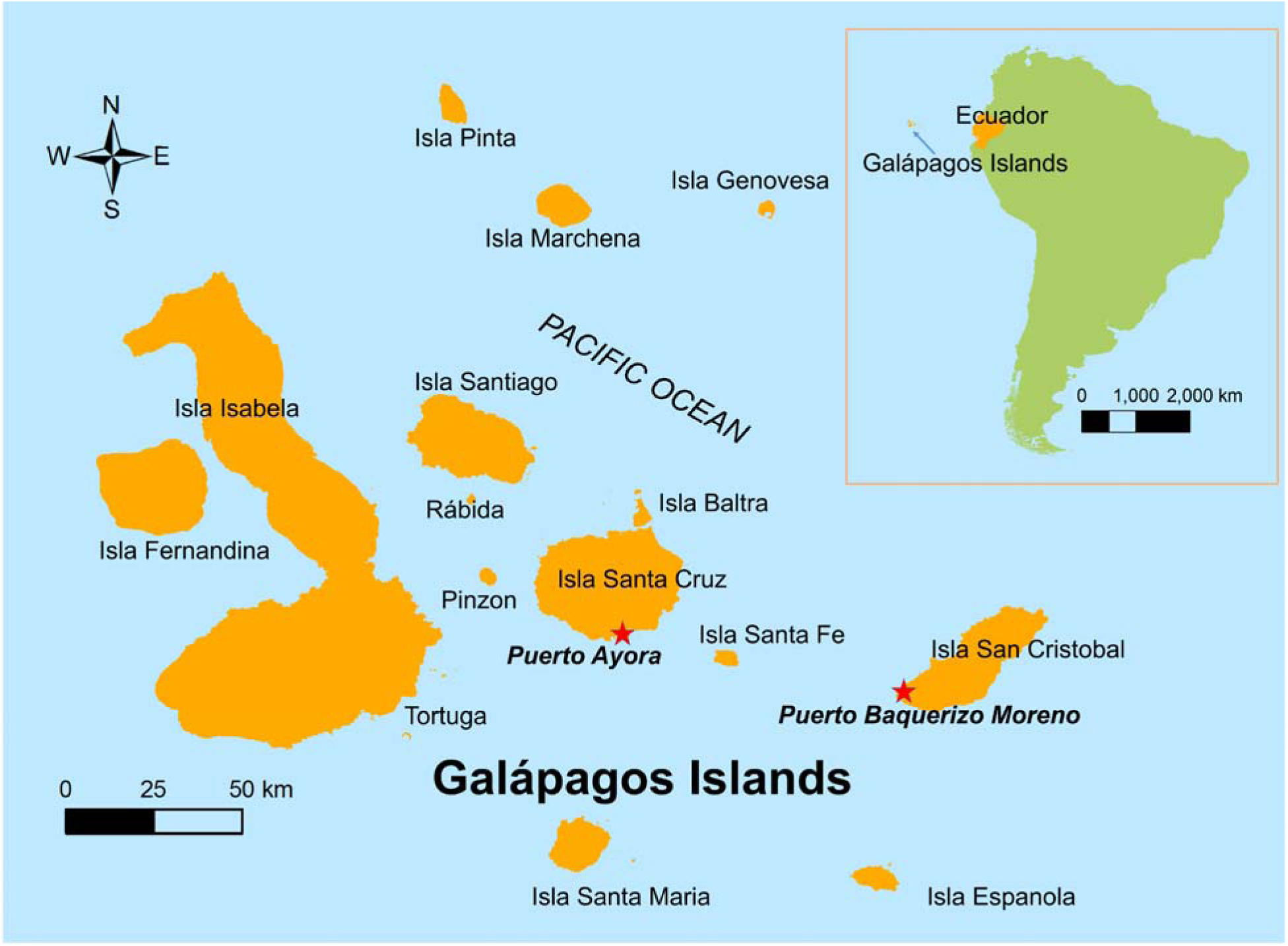
Location of study sites (Puerto Ayora and Puerto Baquerizo) in the Galápagos Islands, Ecuador.

Only three mosquito species have been reported to be present at the Galápagos islands, namely *Aedes aegypti, Aedes taeniorhynchus* and *Culex quinquefasciatus* [16,17] The *Ae. aegypti* mosquito is believed to have been introduced to Santa Cruz Island in the 1990s [17]. The first outbreak of dengue in the Galápagos Islands occurred in 2002 in the city of Puerto Ayora on Santa Cruz Island, the most populated center in the Galápagos (Figures 1 and 2). A total of 227 cases were reported during the outbreak, resulting in an annual incidence rate of 252 cases per 10,000 people. There have been cases of dengue reported from Puerto Ayora every year since then, as well as periodic outbreaks (Figure 2). From 2003 to 2014 a total of 495 cases have been reported, resulting in an average annual incidence rate of 39 cases per 10,000 people. A proportion of cases are thought to be imported from the mainland where dengue is hyper-endemic; however, *Ae. aegypti* is well established and the periodic outbreaks suggest locally acquired infections. Dengue transmission in Puerto Ayora is highly seasonal. Cases peak in June, following the hot season from February to May (23°C − 30°C, mean monthly rainfall = 63 mm; 2002-2014, Figure 3). Puerto Ayora is considered within the dry lowland climatic zone [18].

**Figure 2.**
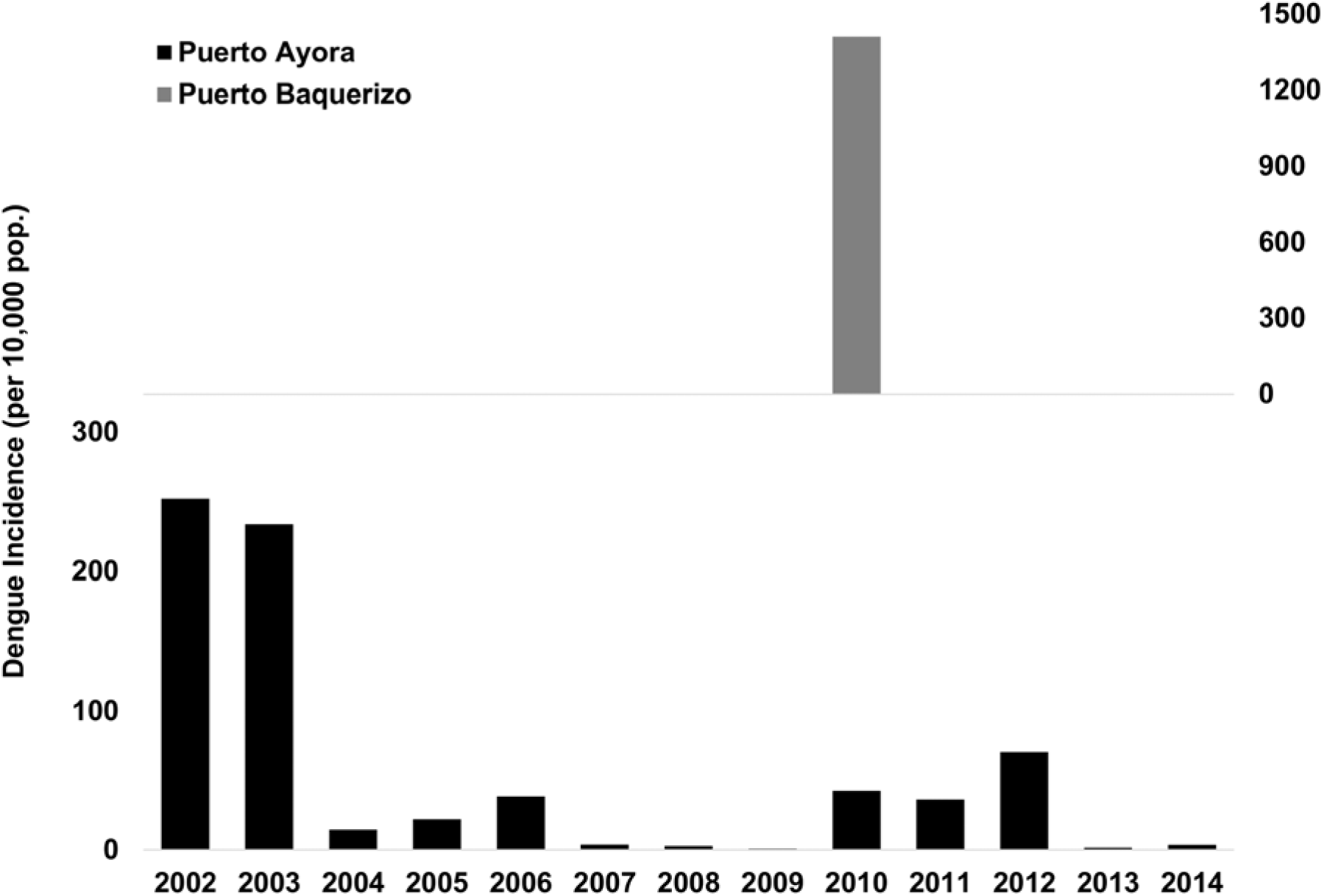
Annual incidence of dengue infections in the Galápagos Islands, showing Puerto Ayora on Santa Cruz Island, and Puerto Baquerizo on San Cristobal Island.

**Figure 3.**
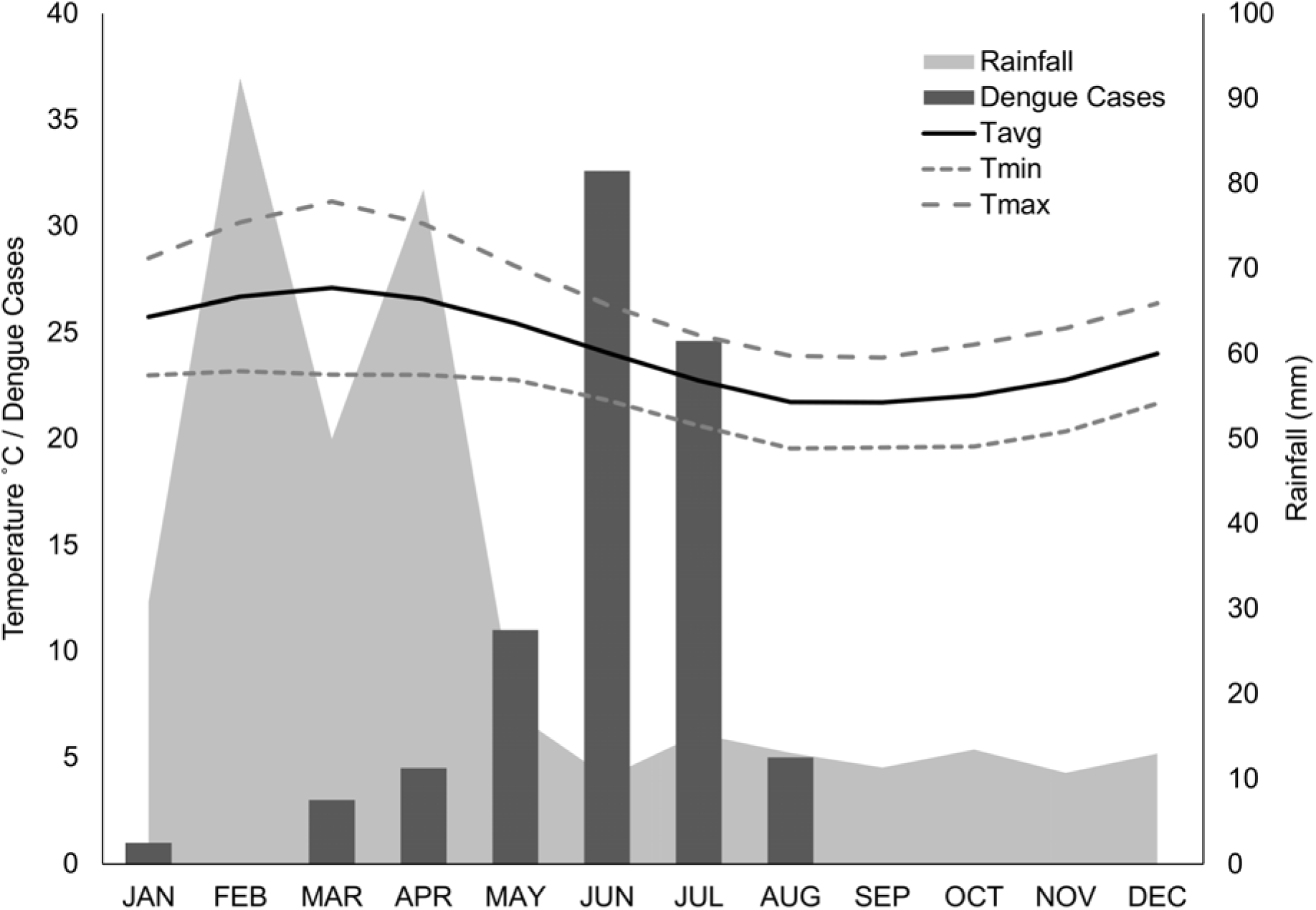
Seasonality of dengue transmission and climate in Puerto Ayora (2000-2012). We show average dengue cases reported by month, average monthly rainfall, and average monthly temperature (Tavg), with average mimimum (Tmin) and maximum (Tmax) monthly temperature. Note that dengue cases appear to peak following the peak of temperature and rainfall, across the same 12 year time period.

San Cristobal, the second-most populated island, experienced a major dengue outbreak in 2010 in the capital city of Puerto Baquerizo Moreno. A total of 941 cases were reported, resulting in an annual incidence rate of 1410 cases per 10,000 people. No cases were reported from 2011 to 2014, the date of this study. Since only one outbreak has been reported, the seasonality of transmission is unknown, although climate conditions are similar to Puerto Ayora.

In Ecuador, dengue is transmitted only by the *Ae. aegypti* mosquito vector, an urbanized anthropophilic mosquito. The dengue virus and its principal mosquito vector, *Aedes aegypti,* were eradicated on mainland Ecuador in the 1950s due to successful DDT campaigns [19]. Following drastic reductions in vector control and rapid, uncontrolled in urbanization in the 1970s and 1980s, dengue re-emerged in Ecuador in 1989; by the early 2000s all four dengue virus serotypes co-circulated in the coastal lowland mainland region [20–22]. The first cases of chikungunya virus (CHIKV - also transmitted by *Ae. aegypti*) were reported in Ecuador at the end of 2014, and a major epidemic emerged in 2015, with over 33,000 cases reported [23]. The first cases of zika virus (ZIKV – also transmitted by *Ae. aegypti*) were confirmed in Ecuador on January 7, 2016. A total of 3,714 suspected and confirmed cases of ZIKV have been reported in Ecuador to date (as of February 16, 2017) [24]. In the Galápagos Islands, suspected dengue cases are confirmed at diagnostic laboratories located in Ministry of Health reference hospitals on each island. A subset of samples are sent on to the virology reference laboratory of the Ministry of Health in Guayaquil, Ecuador. Eighteen confirmed cases of CHIKV in 2015 and two cases of ZIKV in 2016 were reported from the Galápagos Islands [25].

The primary means of preventing dengue transmission in Ecuador is through vector control, reducing the density of *Ae. aegypti* in high-risk households, since a dengue vaccine is not yet available for widespread use. Dengue control is conducted by the Ministry of Health through repeated cycles of ultra-low volume fumigation of neighborhoods throughout the rainy (peak transmission) season; indoor residual spraying in and around homes with suspected dengue cases; routine visits to homes to apply larvicide (temefos/abate) to water-bearing containers, and to destroy larval habitat in and around homes. The central vector control team for the Galápagos is based in Puerto Ayora, due to the greater burden of disease, and a smaller team is based in Puerto Baquerizo. Household-level *Ae. aegypti* control interventions require significant investment of resources including financial resources, personnel, field transportation, chemicals and material supplies. The Ministry of Health in the Galápagos has engaged in multisectoral collaborations for dengue prevention, including dengue education in local schools in partnership with the Ministry of Education, and community clean-up campaigns with private institutions and non-profit organizations.

The aim of our study was to identify the socio-ecological factors that were associated with an increased risk of dengue fever transmission and *Ae. aegypti* presence in households of Puerto Ayora and Puerto Baquerizo Moreno of the Galápagos Islands. To our knowledge, this is the first report on dengue fever risk in the Galápagos Islands. Prior studies in other Latin American locations have shown that risk factors such as water storage practices, knowledge and risk perception, human movement patterns, and housing conditions influence vector abundance and risk of dengue infection [26–30]. Identifying household-level risk factors for dengue transmission in the Galápagos would, therefore, provide specific targets for public health interventions.

## Methods

### Study site

In August and September 2014, we conducted a study in 50 households in Puerto Ayora (PA) (latitude: −0.7402, longitude: −90.31, elevation: 15 m) and 50 households in Puerto Baquerizo (PB) (latitude: −0.9232, longitude: −89.60, elevation: 6 m).). The study was restricted exclusively to urban areas within these two cities. PA and PB are the largest population centers in the Galápagos Islands (2010 population estimates: 11,974 and 6,672, respectively [31]), and the only sites in the Galápagos where cases of autochthonous dengue have been reported. Households were located approximately 200-250 meters apart, the flight range of the *Ae. aegypti* mosquito, in a sector of each city that had historically high *Ae. aegypti* indices according to the local Ministry of Health.

### Field data collection

Trained study technicians surveyed heads of households to identify self-reported prior dengue infections in household members, demographics, risk perceptions, dengue knowledge, sources of dengue information, and vector control practices. Housing conditions were assessed by survey technicians. Survey variables and descriptive statistics are presented in Table 1. The survey instrument was developed from an instrument that has been field tested in other cities in Ecuador [27], and was piloted with Ministry of Health technicians prior to study start (survey instruments in Spanish and English are available upon request).

**Table 1.**
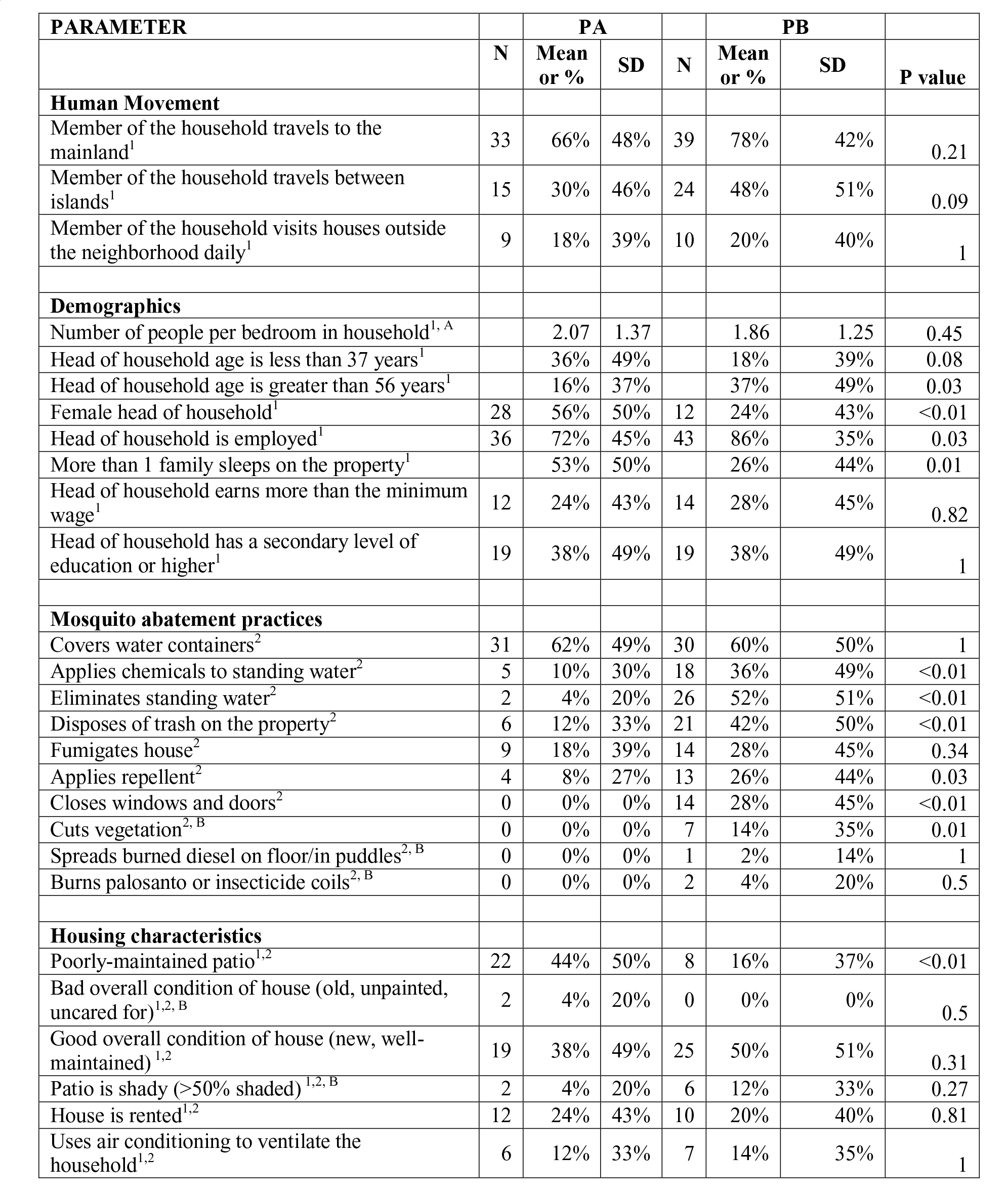
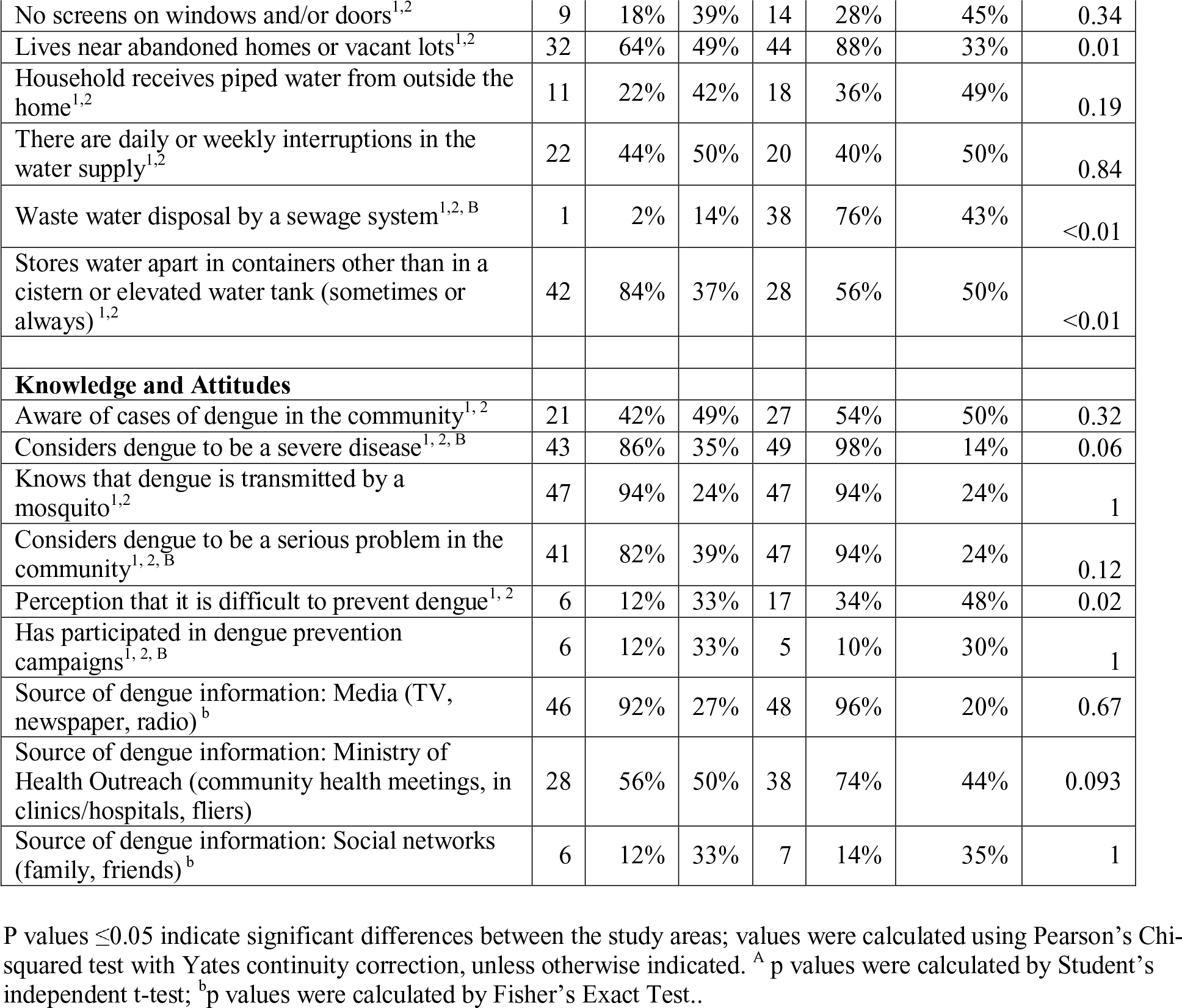
Social-ecological parameters (mean and standard deviation - SD) included in the multimodel selection framework to predict (1) self-reported prior dengue infection and (2) presence/absence of *Aedes aegypti* adults and/or juveniles at the household level for Puerto Ayora (PA) and Puerto Baquerizo Moreno (PB). Model input variables for each model indicated with subscripts (1 = self reported dengue, 2 = *Aedes aegypti*).

| PARAMETER | PA |  |  | PB |  |  |  |
| --- | --- | --- | --- | --- | --- | --- | --- |
|  | N | Mean or % | SD | N | Mean or % | SD | P value |
| **Human Movement** |  |  |  |  |  |  |  |
| Member of the household travels to the mainland<sup>1</sup> | 33 | 66% | 48% | 39 | 78% | 42% | 0.21 |
| Member of the household travels between islands<sup>1</sup> | 15 | 30% | 46% | 24 | 48% | 51% | 0.09 |
| Member of the household visits houses outside the neighborhood daily<sup>1</sup> | 9 | 18% | 39% | 10 | 20% | 40% | 1 |
| **Demographics** |  |  |  |  |  |  |  |
| Number of people per bedroom in household<sup>1, A</sup> |  | 2.07 | 1.37 |  | 1.86 | 1.25 | 0.45 |
| Head of household age is less than 37 years<sup>1</sup> |  | 36% | 49% |  | 18% | 39% | 0.08 |
| Head of household age is greater than 56 years<sup>1</sup> |  | 16% | 37% |  | 37% | 49% | 0.03 |
| Female head of household<sup>1</sup> | 28 | 56% | 50% | 12 | 24% | 43% | <0.01 |
| Head of household is employed<sup>1</sup> | 36 | 72% | 45% | 43 | 86% | 35% | 0.03 |
| More than 1 family sleeps on the property<sup>1</sup> |  | 53% | 50% |  | 26% | 44% | 0.01 |
| Head of household earns more than the minimum wage<sup>1</sup> | 12 | 24% | 43% | 14 | 28% | 45% | 0.82 |
| Head of household has a secondary level of education or higher<sup>1</sup> | 19 | 38% | 49% | 19 | 38% | 49% | 1 |
| **Mosquito abatement practices** |  |  |  |  |  |  |  |
| Covers water containers<sup>2</sup> | 31 | 62% | 49% | 30 | 60% | 50% | 1 |
| Applies chemicals to standing water<sup>2</sup> | 5 | 10% | 30% | 18 | 36% | 49% | <0.01 |
| Eliminates standing water<sup>2</sup> | 2 | 4% | 20% | 26 | 52% | 51% | <0.01 |
| Disposes of trash on the property<sup>2</sup> | 6 | 12% | 33% | 21 | 42% | 50% | <0.01 |
| Fumigates house<sup>2</sup> | 9 | 18% | 39% | 14 | 28% | 45% | 0.34 |
| Applies repellent<sup>2</sup> | 4 | 8% | 27% | 13 | 26% | 44% | 0.03 |
| Closes windows and doors<sup>2</sup> | 0 | 0% | 0% | 14 | 28% | 45% | <0.01 |
| Cuts vegetation<sup>2, B</sup> | 0 | 0% | 0% | 7 | 14% | 35% | 0.01 |
| Spreads burned diesel on floor/in puddles<sup>2, B</sup> | 0 | 0% | 0% | 1 | 2% | 14% | 1 |
| Burns palosanto or insecticide coils<sup>2, B</sup> | 0 | 0% | 0% | 2 | 4% | 20% | 0.5 |
| **Housing characteristics** |  |  |  |  |  |  |  |
| Poorly-maintained patio<sup>1,2</sup> | 22 | 44% | 50% | 8 | 16% | 37% | <0.01 |
| Bad overall condition of house (old, unpainted, uncared for)<sup>1,2, B</sup> | 2 | 4% | 20% | 0 | 0% | 0% | 0.5 |
| Good overall condition of house (new, well-maintained)<sup>1,2</sup> | 19 | 38% | 49% | 25 | 50% | 51% | 0.31 |
| Patio is shady (>50% shaded)<sup>1,2, B</sup> | 2 | 4% | 20% | 6 | 12% | 33% | 0.27 |
| House is rented<sup>1,2</sup> | 12 | 24% | 43% | 10 | 20% | 40% | 0.81 |
| Uses air conditioning to ventilate the household<sup>1,2</sup> | 6 | 12% | 33% | 7 | 14% | 35% | 1 |
| No screens on windows and/or doors<sup>1,2</sup> | 9 | 18% | 39% | 14 | 28% | 45% | 0.34 |
| Lives near abandoned homes or vacant lots<sup>1,2</sup> | 32 | 64% | 49% | 44 | 88% | 33% | 0.01 |
| Household receives piped water from outside the home<sup>1,2</sup> | 11 | 22% | 42% | 18 | 36% | 49% | 0.19 |
| There are daily or weekly interruptions in the water supply<sup>1,2</sup> | 22 | 44% | 50% | 20 | 40% | 50% | 0.84 |
| Waste water disposal by a sewage system<sup>1,2, B</sup> | 1 | 2% | 14% | 38 | 76% | 43% | <0.01 |
| Stores water apart in containers other than in a cistern or elevated water tank (sometimes or always)<sup>1,2</sup> | 42 | 84% | 37% | 28 | 56% | 50% | <0.01 |
| **Knowledge and Attitudes** |  |  |  |  |  |  |  |
| Aware of cases of dengue in the community<sup>1, 2</sup> | 21 | 42% | 49% | 27 | 54% | 50% | 0.32 |
| Considers dengue to be a severe disease<sup>1, 2, B</sup> | 43 | 86% | 35% | 49 | 98% | 14% | 0.06 |
| Knows that dengue is transmitted by a mosquito<sup>1,2</sup> | 47 | 94% | 24% | 47 | 94% | 24% | 1 |
| Considers dengue to be a serious problem in the community<sup>1, 2, B</sup> | 41 | 82% | 39% | 47 | 94% | 24% | 0.12 |
| Perception that it is difficult to prevent dengue<sup>1, 2</sup> | 6 | 12% | 33% | 17 | 34% | 48% | 0.02 |
| Has participated in dengue prevention campaigns<sup>1, 2, B</sup> | 6 | 12% | 33% | 5 | 10% | 30% | 1 |
| Source of dengue information: Media (TV, newspaper, radio)<sup>b</sup> | 46 | 92% | 27% | 48 | 96% | 20% | 0.67 |
| Source of dengue information: Ministry of Health Outreach (community health meetings, in clinics/hospitals, fliers) | 28 | 56% | 50% | 38 | 74% | 44% | 0.093 |
| Source of dengue information: Social networks (family, friends)<sup>b</sup> | 6 | 12% | 33% | 7 | 14% | 35% | 1 |

P values ≤0.05 indicate significant differences between the study areas; values were calculated using Pearson's Chi-squared test with Yates continuity correction, unless otherwise indicated. <sup>A</sup> p values were calculated by Student's independent t-test; <sup>b</sup> p values were calculated by Fisher's Exact Test..

All adult mosquitoes were collected inside and outside of the household using prokopacks, lightweight backpack aspirators shown to be highly effective [32]. Adult mosquitoes were identified to species using a stereo microscope. We conducted standard container surveys to identify the prevalence of water-bearing containers with *Ae. aegypti* pupae and larvae in and around the home [27,33−35]. We recorded descriptive information about each container, including type, use, source of water, and location inside or outside the home. All pupae and a sample of larvae were reared to adults in the laboratory to confirm species identification.

### Statistical models

Survey data were used to identify social-ecological variables associated with the occurrence of self-reported dengue fever and vector presence. We hypothesized that both self-reported dengue and mosquito presence were associated with one or more of these factors (see Table 1). Survey responses were coded and grouped into suites of variables: Human Movement, Demographics, Mosquito Abatement Practices, Housing Characteristics, Knowledge and Attitudes. We used descriptive statistics to compare responses between the two sites, PA and PB, including Student’s independent t-test, Pearson’s Chi-squared test with Yates continuity correction, and Fisher’s Exact Test. All statistics were conducted in R [36].

We used an information theoretical approach to derive best-fit models comprising explanatory variables for 1. self-reported dengue (model: *dengue*) and 2. Presence of *Ae. aegypti* (juveniles and adults, together) at households (model: *mosquito presence*). Two model selection processes were conducted using ‘glmulti’, an R package for multimodel selection, specifying a logistic modeling distribution in a Generalized Linear Model (GLM) framework (GLM, family=binomial, link=logit). The multimodel selection was to determine which survey outcomes related to Human Movement, Demography, Housing Characteristics, and Knowledge and Attitudes were associated with the number of self-reported dengue cases on both islands (Table 1). The second model selection process examined which survey factors related to Housing Conditions, Knowledge and Attitudes, and Mosquito Abatement Practices were influencing the presence of *Ae. aegypti* within the homes of survey participants (Table 1).

Some factors collected in the initial survey were excluded from model variable candidates due to missing or uninformative data (*i.e.,* the responses were identical across households). Selection was run to convergence using glmulti’s genetic algorithm (GA); models were ranked using Akaike’s Information Criterion (AIC) corrected for small sample size (AICc). For each suite of variables in our hypotheses, a best model was obtained (Tables 3 and 4), along with multiple competing top models, using the threshold criteria of AICc ≤ 2 (competing top models given in Suppl. Tables 1, 2). Parameter estimates, odds ratios (OR), and 95% confidence intervals (CI) were calculated for variables in the top ranked model from each search. Variance inflation factors (VIF) and condition numbers (**κ**) were calculated for each top model to assess multi-collinearity and model stability, respectively, with VIF value below 10 indicating low multicollinearity and condition numbers below 30 indicating model stability.

**Table 2.** *Aedes aegypti* (AA) indices in August-September 2014 for households on Puerto Ayora (PA) and Puerto Baquerizo Moreno (PB).

|  | Total | PA |  | PB |  |  |
| --- | --- | --- | --- | --- | --- | --- |
| Vector indices | N | N | % | N | % | P-value |
| Houses inspected | 100 | 50 |  | 50 |  |  |
| Total containers with water | 248 | 119 |  | 129 |  |  |
| Houses with adult AA | 9 | 2 | 4.00% | 7 | 14.00% | 0.16 |
| Houses with containers with juvenile AA | 13 | 3 | 6.00% | 10 | 20.00% | 0.012 |
| Containers with juvenile AA | 16 | 3 | 2.50% | 13 | 10.10% | 0.019 |
| Breteau Index |  | 6 |  | 26 |  |  |
| House Index |  | 6 |  | 20 |  |  |

P-value $\leq 0.05$ indicate significant differences between the study areas; values were calculated using Fisher's Exact Test.

**Table 3.** Summary of the top model for self-reported prior dengue infections.

| Factor | Estimate | St. Error | P-Value | OR (95% CI) | VIF |
| --- | --- | --- | --- | --- | --- |
| Intercept | -12.44 | 3.62 | < 0.001 | . | . |
| People per bedroom | 1.83 | 0.58 | 0.002 | 6.23 (2.42-26.17) | 4.44 |
| Female head of household | -1.91 | 1.08 | 0.077 | 0.15 (0.01-1.05) | 1.59 |
| Head of household earns more than minimum wage | 3.00 | 1.04 | 0.004 | 20.16 (3.15-212.80) | 1.67 |
| Head of household has secondary education of greater | 2.25 | 1.13 | 0.046 | 9.53 (1.28-124.10) | 1.89 |
| House in good condition | 1.74 | 1.10 | 0.114 | 5.71 (0.73-62.26) | 1.95 |
| Visit other neighborhoods daily | -3.47 | 1.80 | 0.054 | 0.03 (0.00-0.51) | 1.55 |
| Travel to continent | 2.32 | 1.19 | 0.051 | 10.14 (1.29-157.45) | 2.04 |
| Travel to island | 3.20 | 1.22 | 0.009 | 24.62 (3.20-454.15) | 2.44 |
| Daily or weekly interruption in the piped water supply | 2.55 | 1.14 | 0.026 | 12.75 (1.77-176.58) | 2.13 |
| No screens on windows or doors | -3.18 | 1.58 | 0.044 | 0.04 (0.00-0.55) | 2.11 |
| Aware of cases of dengue in the neighborhood | 2.54 | 1.16 | 0.028 | 12.62 (1.81-202.17) | 2.05 |

\* AICc=69.89, $\kappa$ = 12.97

**Table 4.** Summary of the top model for the presence of adult and juvenile *Aedes aegypti.*

| Factor | Estimate | St. Error | P-Value | OR (95% CI) | VIF |
| --- | --- | --- | --- | --- | --- |
| Intercept | -3.33 | 2.42 | 0.019 | . | . |
| Uses mosquito nets | 3.40 | 1.22 | 0.005 | 29.86 (3.57-488.97) | 2.46 |
| Covers water containers | -2.85 | 0.97 | 0.003 | 0.05 (0.01-0.32) | 1.53 |
| Patio in bad condition | 1.93 | 1.01 | 0.056 | 6.89 (1.03-60.93) | 1.55 |
| House in bad condition | -19.87 | 2562.01 | 0.994 | < 0.001 (NA) | 1.00 |
| Air conditioning | 2.93 | 1.05 | 0.005 | 18.76 (2.75-192.55) | 1.46 |
| No screens | 3.18 | 1.31 | 0.015 | 23.95 (2.41-474.23) | 2.23 |
| Dengue is difficult to prevent | 3.29 | 1.04 | 0.002 | 26.90 (4.47-308.79) | 1.77 |
| Dengue is a problem | -1.94 | 1.53 | 0.206 | 0.14 (0.01-2.77) | 1.70 |

\* AICc=63.00, $\kappa$ = 12.41

## Results

### Dengue infections and Ae. aegypti abundance

Respondents from 100 households were interviewed, with PA households (n = 50) representing 78 household members, and PB households (n = 50) representing 152 household members. In PA, 11/77 (14.3%) of people reported prior dengue infections. In PB, 37/150 people reported prior dengue infections (24.7%). At the household level, prior dengue infections were reported by more households on PB (24.3%) than on PA (14.1%), although the difference was not significant (p>0.05, Chi-squared test). Most people on both islands reported seeking medical care when ill with dengue (PA = 64%, PB = 78%, p>0.05, Fisher’s Exact Test).

A total of 248 water-bearing containers were inspected for the presence of *Ae. aegypti* (PA = 119, PB = 129). Significantly more houses were found to have containers with juvenile *Ae. aegypti* in PB than in PA (p=0.012, Table 2). House Indices (number of homes with juvenile *Ae. aegypti* per 100 homes) in PB were 20 and in PA were 6. Breteau Indices (number of containers with juvenile *Ae. aegypti* per 100 homes) in PB were 26 and in PA were 6. A greater proportion of surveyed containers were found with *Ae. aegypti* juveniles in PB than in PA (p=0.019). The predominant characteristics of containers positive for juvenile *Ae. aegypti* (n=16) were: low water tanks made of cement or plastic (92%); containers that were completely or partially uncovered (92%); containers located outdoors (85%); containers that were shaded (85%); containers filled with tap water as opposed to rain water (100%); and containers intended for domestic use (*i.e.,* used for cooking, cleaning, laundry as opposed to abandoned containers) (77%).

### Risk perceptions and practices

Most households on PA (82%) and PB (94%) reported that dengue was a serious problem in their community (p = 0.1) and a severe disease (PA = 86%, PB = 98%, p = 0.06) (Table 1). Significantly more PB households reported that it was difficult or impossible to prevent dengue (p = 0.02, Table 1). The majority of heads of households knew that dengue was transmitted by a mosquito (94% on both islands), and most people had received information about dengue prevention (64% on both islands). However, few people had participated in dengue prevention campaigns (PA = 12%, PB = 10%). Sources of dengue information were similar between islands, with media (TV, newspaper, radio) as the primary source of information on both islands, and social networks as the least likely source of information (Table 1).

Overall, PB households implemented more prevention strategies than PA households (PA mean=1.48, SD=0.76, PB mean=3.76, SD=2.09, p<0.001). PB households reported several prevention strategies significantly more frequently than PA households, including use of screens on windows and doors (p<0.001), topical repellent (p=0.03), keeping the property clear of trash (p=0.02), closing windows and doors (p<0.001), cutting grass and plants (p=0.01), adding chemicals to standing water (p=0.004), and eliminating standing water (p<0.001) (Table 1). Fumigation and use of mosquito nets were the least commonly reported mosquito control actions.

### Model selection outcomes

The top-ranked model of a prior self-reported case of dengue in the household (AICc=69.89, **κ**= 12.97) included the following suite of positively associated variables: the number of people per room in the home, the head of the household earning more than minimum wage, the head of the household with secondary education or higher, having a house in good condition, household members who travel to the continent, household members who travel between islands, frequent interruptions in the piped water supply, and being aware of dengue cases in their community (Table 3, Figure 4). Having a female head of household, visiting other neighborhoods daily, and no screens on doors and windows were negatively associated with self-reported dengue. Nineteen additional models were found within 2 AICc units of the top model (Supplementary Table 3).

**Figure 4.**
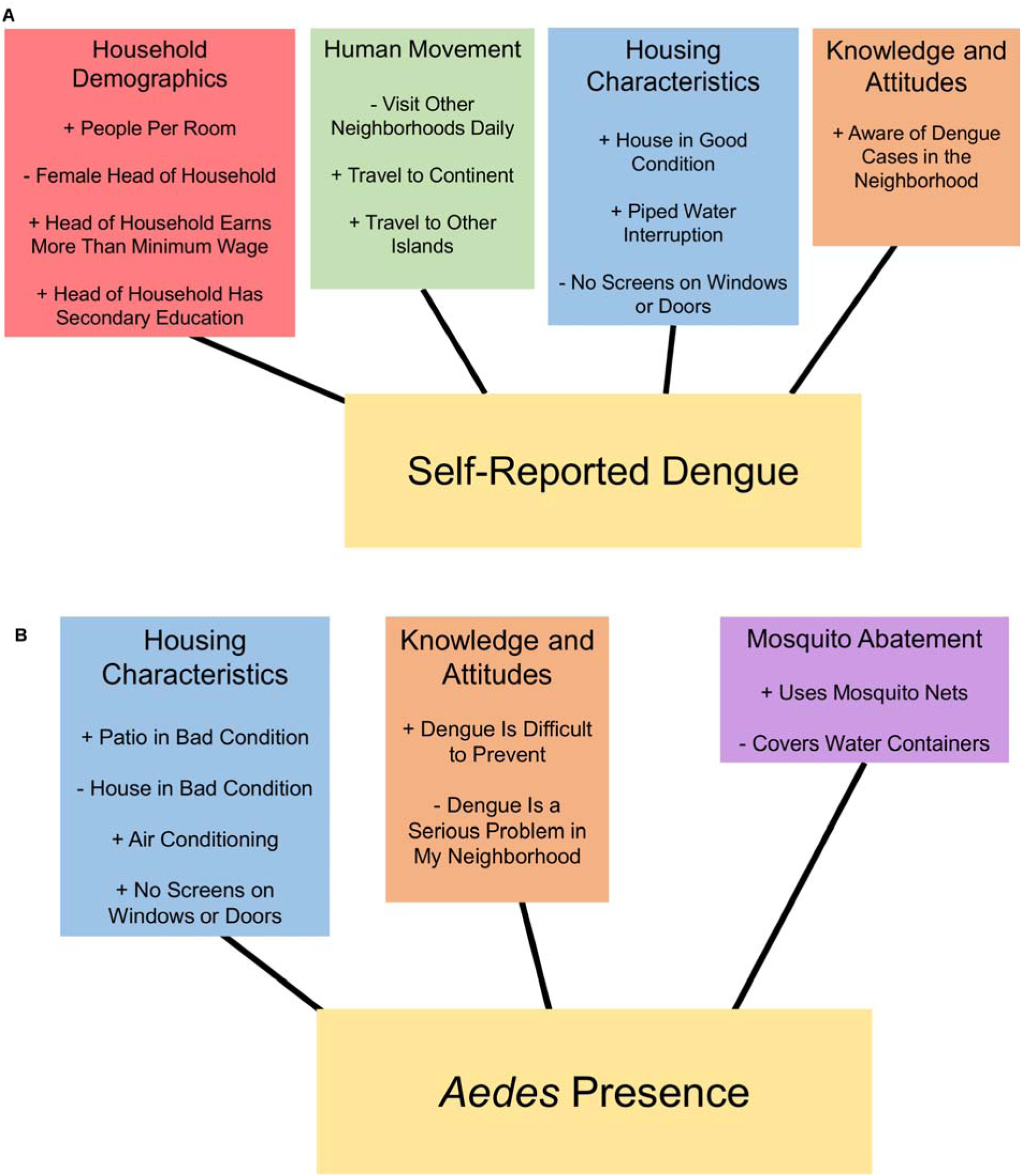
Suites of socio-ecological variables in the top selected models for A. self-reported dengue cases, and B. presence of *Aedes aegypti.*

The top-ranked model to predict the presence of *Ae. aegypti* in households (AICc=63.00, **κ**= 12.41) included the following positively associated variables: use of mosquito nets in the home, bad patio condition, air conditioning, no screening on windows or doors, and the perception that dengue is a difficult disease to prevent (Table 4, Figure 4). Negatively associated variables included covering water containers, bad house condition, and the perception that dengue is a problem. Eight additional models were found within 2 AICc units of the top model, comprising alternating selections of similar variables to the top model (Supplementary Table 4).

## Discussion

The emergence of dengue in the Galápagos Islands, and the associated risk of imported and autochthonous chikungunya and zika infections, have serious implications for the tourism-based economy of the islands. Tourism is an important and substantial source of income for inhabitants of the Galápagos Islands, with 13% of Santa Cruz inhabitants and 7% of San Cristóbal inhabitants reporting tourism as their source of income according to the 2010 census [37]. The risk of disease could discourage tourists from visiting the islands. Furthermore, the high volume of tourists traveling from mainland Ecuador to the islands, particularly tourists passing through the city of Guayaquil, a dengue endemic city, could present a source of reintroduction of the virus.

This study provides the first insights into the nature of household level dengue fever risk and *Ae. aegypti* presence on the Galápagos Islands of Ecuador, where the disease has emerged in the last 15 years. Important factors associated with reported dengue and mosquito presence are described as suites of socioecological factors (Figure 4): human movement, household demographics, housing characteristics, knowledge and attitudes, and mosquito abatement practices. These findings can be readily interpreted and used to inform the design and implementation of targeted vector control campaigns that reflect the local social-ecological context [27,38].

We found that self-reported prior dengue infections were associated with housing conditions, dengue awareness, higher income and education of the head of the household, and frequent travel by household members (Figure 4, Table 3). Higher income and education of the head of the household were positively associated with frequent travel by household members and greater awareness of dengue. The role of human movement in dengue transmission has been documented in prior studies in Iquitos, Peru [28,39].

Greater housing density (people per bedroom) and frequent interruptions in the piped water supply were also indicative of greater dengue risk. These variables likely reflect greater risk of exposure to infectious mosquito bites, due to larval habitat in water storage containers, and increased probability of infectious bites due to human crowding. These findings are consistent with other studies, both on mainland Ecuador [26,27,40], and elsewhere [41–43].

The important variables in the mosquito presence model indicate that housing conditions, dengue risk perception, and prevention practices are important risk factors. We found that homes were more likely to have *Ae. aegypti* if they had no screens on windows or doors, if they did not cover water containers, and if they perceived that dengue was difficult to prevent (Figure 4). These risk factors are consistent with prior studies from Ecuador, Taiwan, and India [27,43,44]. Interventions to address these factors include dengue awareness and community mobilization campaigns [45], water container covers, and programs to provide low-cost screening to homeowners. Paradoxically, we found that the use of mosquito nets was positively correlated with *Ae. aegypti* presence. This may be as simple as a causal reverse in a correlation – bet nets are more likely to be used when mosquitoes are perceptibly present. However, it may instead be the case of the wrong intervention for the vector. *Ae. aegypti* have a small range and will bite during the day (in contrast to other mosquito genera such as *Anopheles*), so bed nets may not be an effective barrier to dengue transmission. Another unexpected finding was that air conditioning was positively associated with *Ae. aegypti* presence. This result was counterintuitive, because one would expect homes with air conditioning to have closed windows and subsequently fewer mosquitoes and lower dengue risk, as shown in prior studies [46]. However, it is possible that water buildup and puddles created by air conditioning units could create mosquito habitat, and this requires further investigation, including understanding the context in which air conditioning units are installed, versus used. If rooms are cooled for a few hours a day only, and in the evening, windows are used to cool houses, an air conditioning unit is a false signal of closed window behavior. A more comprehensive study should include AC practices in questionnaires.

Our findings highlight the importance of human movement in determining dengue transmission, particularly on islands where people travel regularly to the mainland, which can lead to frequent reintroduction of the virus and its vector. Individuals with frequent travel to other islands, as well as between the islands and the continent, may be considered to be at higher risk for dengue infection. Previous studies of dengue on islands have emphasized the importance of preventing the transmission of dengue from the mainland [47], as well as reintroduction from surrounding islands [48]. Prior studies of West Nile Virus risk in the Galápagos found that the transportation of infected mosquitoes via airplanes was the most likely means that the virus would be introduced to the islands [13]. The transmission dynamics between mainland and island populations may be exacerbated by seasonal differences in the epidemiology of travelers [49], which may support the restriction of travelers during significant outbreaks on the mainland. The recent emergence of chikungunya and zika viruses highlights the importance of understanding these regional movement and transmission dynamics [50,51]. Our findings also suggest the importance of local, community-based movement dynamics in the transmission dynamics of dengue, as has been explored in previous studies [28].

Water storage and access were among the most important household risk factors for both prior dengue infections and the presence of *Ae. aegypti,* as shown in prior studies in mainland Ecuador [27,40]. Water access is a serious concern on the Galápagos Islands, which has a limited supply of fresh water [52,53]. As a result, many inhabitants store water around the home for daily use [54], creating the ideal habitat for *Ae. aegpyti* juveniles, especially in PB. The characteristics of containers positive for juvenile *Ae. aegypti* (uncovered water storage containers located outdoors) indicates that community clean-up campaigns focused on the elimination of rubbish in the patio may have a limited effect on *Ae. aegypti* abundance, at least during the cool season, when this study was conducted. The primary focus in these areas should be the creation of sealed (*Aedes-*proof) water storage containers used by households and located outdoors in the patio. This result highlights the importance of integrated household water management strategies in regions that are water-scarce and at risk of dengue risk.

We found that there were differences in self-reported prior dengue infections, vector abundance, prevention strategies, sources of information, and risk perception between PA and PB. This suggests that there may be differences in a number of factors between the islands, including disease burden, community outreach programs, community awareness, and/or access to information between the two sites. The high proportion of homes with juvenile *Ae. aegypti* on PB indicate that there was significant risk of another dengue outbreak, even during the low transmission season when this study was conducted.

PB households that reported prior dengue infections were most likely from the dengue outbreak that occurred in 2010 (four year prior), when most of the population was susceptible to dengue infections. The prevalence of past dengue infections reported by surveyed households in this study (24.6%) was higher than the Ministry of Health reported dengue prevalence in 2010 (14.1%). The burden of disease during the outbreak was likely higher than reported by the Ministry of Health, since subclinical and mild infections are not captured by passive surveillance systems [55]. This discrepancy may also indicate that people are aware of the symptoms but did not report those symptoms to authorities; and we see considerable self-diagnosed, but unreported dengue. A dengue surveillance study conducted in the same year in southern coastal Ecuador found that 44% of people with acute or recent dengue infections reported no dengue-like symptoms, and there were an additional three dengue infections in the community for each case reported by the Ministry of Health [56]. This discrepancy is worth exploring in future work, as it indicates higher potential rates of infection than is informing current policies.

The results of this study indicate that very few households have been actively engaged in dengue control campaigns, despite the perceived importance and high awareness of the disease. A survey conducted in PA and PB in the same year found that nine out of ten people felt that they were not prepared for a future dengue outbreak, and about a third of people reported that they were vulnerable to dengue infections (F. Ortega,*pers. comm*.). Studies from mainland Ecuador identified factors that influenced people’s willingness to engage in vector control, including social cohesion and leadership, the perceived role of the government versus the community, and the time and financial costs of household vector control [26,57]. An integrated program of vector control and environmental interventions on the islands has focused on the elimination of used tires (potential *Ae. aegypti* larval habitat), removing them from the islands to the continent to be recycled and converted into road cover and turf for athletic fields, among other uses. Tires were also used to waterproof landfills on the islands. From 2012 to 2014, 35,000 tires were removed from the islands to the continent. This was a coordinated intervention between the Ministry of Health and the Ministry of Environment, and recommended by PAHO [58,59]. These strategies complemented traditional vector control interventions, and aimed to reduce vector densities on Santa Cruz and San Cristobal, where used tires were completely removed.

Another source of introductions of vectors are cargo ships that carry food and other products for consumption by the Galápagos population and tourists. Invasive species are an ongoing risk for the islands, and vector introduction could be avoided by strengthening vector surveillance on cargo ships.

Future studies that explore barriers to dengue control could be used to inform the development of community-based interventions to reduce the risk of dengue and other *Ae. aegypti* transmitted diseases, through strategies such as Communication for Behavioral Impact (COMBI) [45,60,61], which is supported by the Pan American Health Organization (PAHO) and the Ecuadorian government. In 2006, a COMBI project, supported by PAHO and the U.S. Centers for Disease Control (CDC), was implemented on Santa Cruz Island with the goal of cleaning and scrubbing water storage containers to reduce the density of *Ae. aegypti* [60]. In this multi-sectoral project, high school students successfully encouraged households to cleaned their water tanks [62]; however, the sustainability of the project was limited by the reliance on students to conduct the household inspections and limited engagement of householders [60].

One of the greatest public health challenges observed during this study was the implementation of uniform vector control and surveillance across the four populated islands, which are more than two hours apart by boat. The higher larval indices on PB may be due to less staffing and resources for vector control, since there have been fewer total cases reported on PB than PA. Investigating the sources of these discrepancies could provide insight on how to better mobilize and engage communities in order to promote the adoption of preventative behaviors across relatively isolated islands at risk of emerging infectious diseases.

## Conclusions

To our knowledge, this is the first study of dengue risk and *Ae. aegypti* in the Galápagos Islands. The findings that human movement within and between islands, and to and from the mainland, were important to reported dengue cases, confirms concerns of this route of introduction and repeated transmission. Bolstering surveillance of both tourism and cargo routes of entry would be useful to mitigate potential further introductions. The identification of both sources of knowledge and perceptions, and our assessment of both the importance of knowledge of prevention, and a pervasive lack of involvement in control campaigns, point to targets for policy and action. We found that, similarly to studies conducted in mainland Ecuador, housing condition, and water supply, access, and storage related behaviors are important. The water connection is particularly poignant for the Galápagos, where access to freshwater is a concern. Given the geographic challenges faced in distributed island vector management, this is a complicated and unique setting for dengue management, but similarities with other studies, in terms of targets for interventions identified in this study, provide useful information for establishing combined outreach and direct intervention efforts in the public health arena.

## Declarations

### Ethics approval and consent to participate

The study protocol was reviewed and approved by the Institutional Review Board (IRB) of the Universidad San Francisco de Quito and SUNY Upstate Medical University. The protocol was also approved by the Agencia de Regulación y Control de la Bioseguridad y Cuarentena para Galápagos (ABG) prior to study start. Heads of households (>18 years of age) were consented by trained study technicians and signed an informed consent form prior to study start.

### Consent for publication

Not applicable

### Availability of data and material

The datasets used and/or analyzed during the current study are available from the corresponding author on reasonable request.

## Author contributions

AMSI, MBC, RL, and FO were responsible for initial conception of the study, and AMSI was responsible for the design of the field study. AMSI, EG, RL and MC were responsible for data collection. RN, AMSI CL, MBC, and SR were responsible for data analysis and interpretation. RN, AMSI, and SR were responsible for drafting the article. All authors were responsible for critical revision and final approval of the version to be published.

## Acknowledgments

Thank you to the community members from Puerto Ayora and Puerto Baquerizo Moreno for their participation in this study. In particular, thank you to Dr. David Basantes field technicians from the Ministry of Health and the Agencia de Regulación y Control de la Bioseguridad y Cuarentena para Galápagos (ABG).

---

**Supplementary Table 3:**
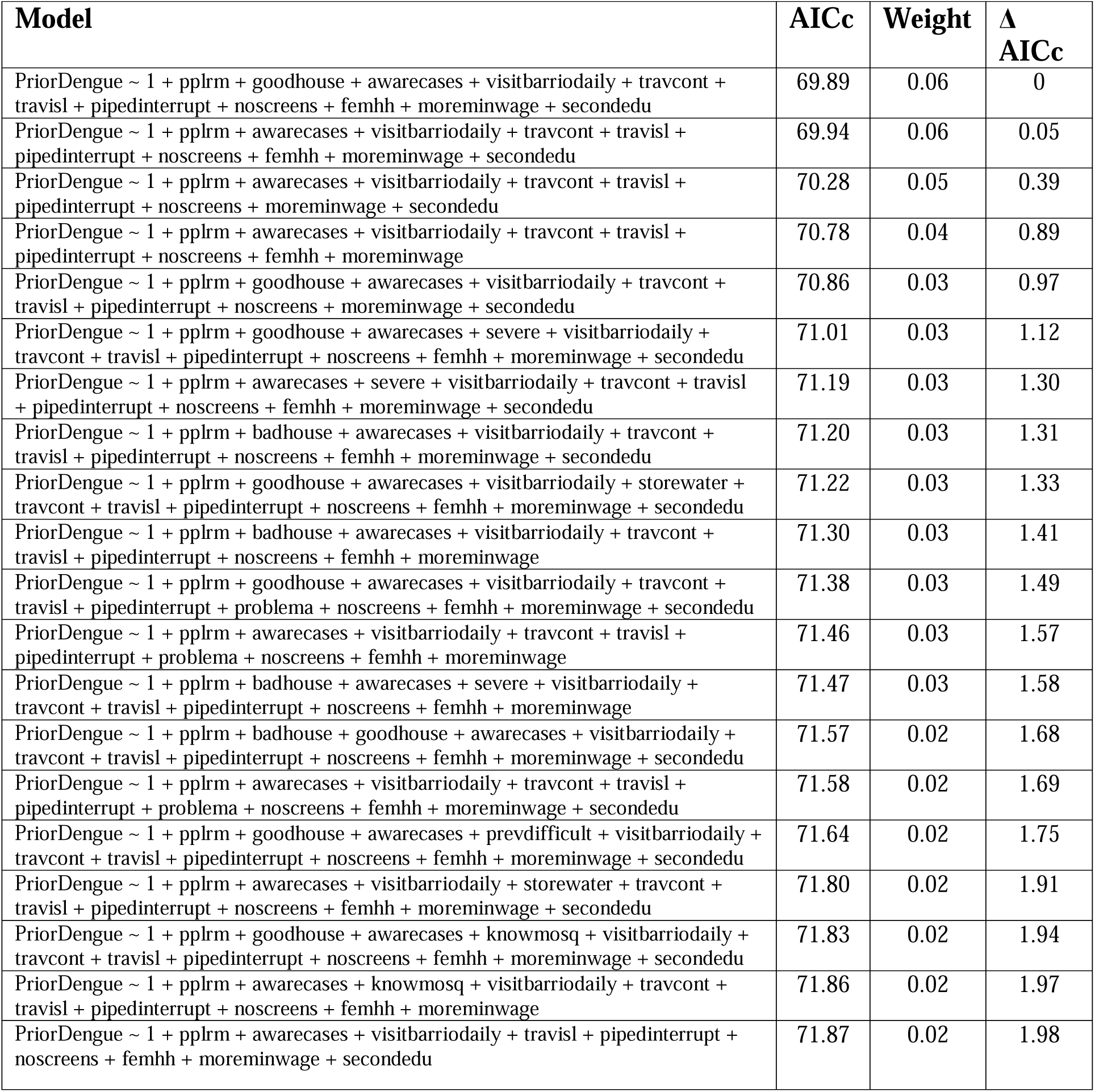
Top models within 2 AICc units for self-reported dengue model.

**Supplementary Table 4:** Top models within 2 AICc units for *Aedes aegypti* presence.

| Model | AICc | Weight | $\Delta$ AICc |
| --- | --- | --- | --- |
| AdultCont_pres ~ 1 + mallas + tapar + badpatio + badhouse + prevdifficult + AC + noscreens | 63.00 | 0.08 | 0 |
| AdultCont_pres ~ 1 + mallas + tapar + badpatio + badhouse + prevdifficult + AC + problema + noscreens | 63.74 | 0.06 | 0.75 |
| AdultCont_pres ~ 1 + mallas + tapar + badhouse + prevdifficult + AC + noscreens | 63.96 | 0.05 | 0.96 |
| AdultCont_pres ~ 1 + mallas + tapar + badpatio + badhouse + severe + prevdifficult + AC + noscreens | 64.18 | 0.05 | 1.18 |
| AdultCont_pres ~ 1 + mallas + tapar + badpatio + badhouse + prevdifficult + AC + pipedout + noscreens | 64.51 | 0.04 | 1.51 |
| AdultCont_pres ~ 1 + mallas + tapar + cerrar + badpatio + badhouse + prevdifficult + AC + noscreens | 64.55 | 0.04 | 1.55 |
| AdultCont_pres ~ 1 + mallas + tapar + badpatio + badhouse + prevdifficult + abandon + AC + noscreens | 64.59 | 0.04 | 1.59 |
| AdultCont_pres ~ 1 + mallas + tapar + badpatio + badhouse + prevdifficult + AC + storewater + noscreens | 64.91 | 0.03 | 1.91 |
| AdultCont_pres ~ 1 + mallas + tapar + diesel + badpatio + badhouse + prevdifficult + AC + noscreens | 64.99 | 0.03 | 1.99 |

